# Data-independent acquisition protease-multiplexing enables increased proteome sequence coverage across multiple fragmentation modes

**DOI:** 10.1101/2021.07.15.452565

**Authors:** Alicia L. Richards, Kuei-Ho Chen, Damien B. Wilburn, Erica Stevenson, Benjamin J. Polacco, Brian C. Searle, Danielle L. Swaney

## Abstract

The use of multiple proteases has been shown to increase protein sequence coverage in proteomics experiments, however due to the additional analysis time required, it has not been widely adapted in routine data-dependent acquisition (DDA) proteomic workflows. Alternatively, data-independent acquisition (DIA) has the potential to analyze multiplexed samples from different protease digests, but has been primarily optimized for fragmenting tryptic peptides. Here we evaluate a DIA multiplexing approach that combines three proteolytic digests (Trypsin, AspN, and GluC) into a single sample. We first optimize data acquisition conditions for each protease individually with both the canonical DIA fragmentation mode (beam type CID), as well as resonance excitation CID, to determine optimal consensus conditions across proteases. Next, we demonstrate that application of these conditions to a protease-multiplexed sample of human peptides results in similar protein identifications and quantitative performance as compared to trypsin alone, but enables up to a 63% increase in peptide detections, and a 27% increase non-redundant amino acid detections. Importantly, this resulted in 100% sequence coverage for numerous proteins, suggesting the utility of this approach in applications where sequence coverage is critical, such as proteoform analysis.

## Introduction

Comprehensive detection and quantification of the proteome is essential to enhancing our understanding of human biology and disease. Beyond advances in data acquisition and instrumentation, mass spectrometry (MS) based efforts to maximize proteome coverage have historically focused on increasingly extensive fractionation of complex tryptic peptide mixtures prior to mass spectrometric analysis, pushing the number of detectable proteins to a level comparable to modern transcriptomics (∼14,000 proteins).^1–3^ Despite such advances in protein detection, an inherent limitation exists in our ability to more completely characterize amino acid sequence coverage due to the near exclusive use of trypsin for proteolytic cleavage during sample preparation. Trypsin is a well suited protease for proteomics, as it produces peptides with a basic residue (Arg or Lys) on the C-terminus, and when protonated in the gas-phase, these peptide cations are ideal for sequencing by collisional-activation tandem MS (i.e., CID MS/MS).^4,5^ However, it also locks a considerable fraction of the proteome in peptides either too small or too large for routine MS identification, thus rendering these regions of the proteome effectively invisible in nearly all proteomics experiments performed to date.^6^ Undoubtedly, these hidden regions contain a wealth of undiscovered and important biological knowledge regarding protein isoforms, alternative splicing events, and post-translational modification (PTMs).

A straightforward method to uncover these hidden regions is by the use of additional proteases, which enables individual amino acids to be represented by alternative peptides sequences.^1,6–16^ Basic residues are not uniformly distributed across the proteome, and a prime example of this is the analysis of histone tails where the PTM status regulates chromatin structure and gene expression. Histone tails are challenging to analyze with trypsin due to the high density of basic amino acids,^17^ and thus digestion with alternative proteases (e.g. GluC) has been essential to deciphering the combinatorial code of histone PTMs.^18^ Furthermore, the use of alternative protease has been demonstrated to substantially increase the detection of both nonsynonymous single nucleotide variants (nsSNVs)^19^ and phosphorylation sites.^20^

Despite the demonstrated utility of using non-tryptic proteases, they have not been widely adopted into the standard proteomics workflow. Using standard data-dependent acquisition (DDA) MS methods, where precursor peptides are individually isolated and fragmented according to precursor intensity, samples generated from each protease must be analyzed independently. Consequently, the inclusion of additional proteases dramatically increases the amount of instrument time required per experiment. Additionally, there is a diminishing return on investment, as non-tryptic proteases generate peptides with less ideal characteristics for CID fragmentation,^21^ resulting in fewer peptide identifications per sample as compared to trypsin. Combined, these factors have resulted in the limited usage of non-tryptic proteases to specific biological applications.

Over the past decade, data-independent acquisition (DIA) has matured as an alternative acquisition strategy to DDA. DIA methods sequentially sweep across m/z precursor isolation windows to acquire tandem mass spectra irrespective of which peptides are being sampled.^22^,23 This approach eliminates the requirement that at least one MS/MS is collected for each peptide, and opts instead to acquire MS/MS that intentionally contain multiple peptides. While deconvolving peptide signals from this type of data can be more challenging, the use of peptide-centric^24^ searching has significantly improved peptide detection rates. This new paradigm presents the opportunity to multiplex proteomic samples generated from a variety of different proteases in a single DIA-MS analysis,^25^ and consequently significantly increasing proteome sequence coverage while incurring minimal increases in MS instrument time. Furthermore, results by Bruderer et al. demonstrate the ability to detect more peptides in a single LC-MS/MS experiment than the number of acquired MS/MS spectra, and imply a hidden capacitance of DIA to detect more peptides within a given dynamic range.^26^ However, to date DIA has been highly optimized for tryptic peptides fragmented by higher energy beam-type CID (bt-CID), with few reported examples of the use of non-tryptic proteases^25^ or lower energy resonance excitation CID (re-CID).^23,27^ As a result, high resolution peptide libraries suitable for analyzing DIA measurements of non-tryptic peptides are extremely limited, and methods to generate DIA-only libraries^28^ are constrained by lower-accuracy fragmentation prediction for non-tryptic peptides using current modeling algorithms.^29^

Here we evaluate DIA approaches for non-tryptic peptides and then exploit the excess sequencing capacity and library-based detection in DIA to multiplex proteomics samples resulting from a mixture of different proteases to achieve significantly greater information return from a single MS acquisition.

### Experimental Section

#### Sample preparation

HEK 293T, HSC6, and SCC25 cell pellets were resuspended in a lysis buffer composed of 8 M urea, 100 mM ammonium bicarbonate (ABC) and 100 mM NaCl (pH∼8). Cells were lysed via probe sonication, on ice, at 20% amplitude for 20 seconds, followed by 10 seconds rest. In total, this process was performed three times. Lysate protein concentration was measured by Bradford assay. Disulfide bond reduction and carbamidomethylation of cysteines was performed by the addition of a 1:10 volume of reduction/alkylation buffer (100 mM tris(2-carboxyethyl)phosphine (TCEP) and 400 mM 2-chloroacetamide (CAA)) to the lysate. Samples were then incubated for 5 minutes at 45°C with shaking (1,500 rpm). Lysate was diluted to 1.5 M final urea concentration with 100 mM ABC, 100 mM NaCl solution. Lysate was split into three individual samples and digested overnight with either trypsin (Promega), GluC (Worthington Biochem), or AspN (Promega) at 37°C at a protease:substrate ratio of 1:100 (w:w). Following digestion, peptides were acidified with 10% trifluoroacetic acid (TFA) to a pH∼2. Peptides were desalted using C18 tips (Nest Group). Tips were activated with 0.1% TFA/80% acetonitrile (ACN) and equilibrated with 0.1% TFA. Following the addition of peptide samples, tips were washed with 0.1% TFA. Peptides were eluted with 0.25% formic acid (FA)/50% ACN. Samples were dried down by SpeedVac and resuspended in 0.1% FA. To generate digest mixtures, trypsin, AspN, and GluC generated peptides were combined (w:w:w) in the following ratios (trypsin:GluC:AspN) : 1:1:1 (equal amounts of all digest mixtures), 1:2:2 (twice as much GluC and AspN digests as tryptic digest), and 1:3:3 (three times as much GluC and AspN digests as tryptic digest).

#### LC-MS/MS analysis

For HEK 293T digests, reversed phase columns were prepared in house. 75–360 μm inner-outer diameter bare-fused silica capillary with an electrospray tip (New Objective) was packed with 1.7 μm diameter, 130 Å pore size, Bridged Ethylene Hybrid C18 particles (Waters) to a length of 15 cm. The column was installed on a Easy nLC 1200 ultra-high pressure liquid chromatography system (Thermo Fisher Scientific) interfaced via a Nanospray Flex nanoelectrospray source. Mobile phase A consisted of 0.1% FA, and mobile phase B consisted of 0.1% FA/80% ACN. Peptides were separated by an organic gradient from 2% to 28% mobile phase B over 61 minutes followed by an increase to 44% B over 9 minutes, then held at 88% B for 10 minutes at a flow rate of 350 nL/minute. Analytical columns were equilibrated with 6 mL of mobile phase A.

For HSC6 and SCC25 digests, a 15 cm PepMap RSLC column (Thermo Fisher Scientific) packed with 3µm particles was installed on a Easy nLC 1200 ultra-high pressure liquid chromatography system (Thermo Fisher Scientific) interfaced via a nanoelectrospray source. Mobile phase A consisted of 0.1% FA, and mobile phase B consisted of 0.1% FA/80% ACN. Peptides were separated by an organic gradient from 2% to 28% mobile phase B over 93 minutes followed by an increase to 44% B over 17 minutes, then held at 88% B for 10 minutes at a flow rate of 300 nL/minute.

For DDA analysis of HEK 293T (Lumos) and SCC25 and HSC6 (Eclipse) digests, eluting peptide cations were analyzed by electrospray ionization on an Orbitrap Fusion Lumos (Thermo Fisher Scientific). MS1 scans of peptide precursors were performed at 120,000 resolution (200 m/z) over a scan range of 350-1050 m/z, with an AGC target of 250% and a max injection time of 100 ms. MIPS was set to peptide mode, and charge states 2-6 were selected for fragmentation. Dynamic exclusion was set to 20 s with a 10 ppm tolerance around the precursor and isotopes. The instrument was run in Top Speed Mode with a 3 second setting. Tandem MS was performed via quadrupole isolation with a width of 1.4 Th. For re-CID experiments, CID fragmentation was performed at a normalized collision energy (NCE) of either 24, 27, 30, or 33%, CID activation time of 10 ms, and activation Q of 0.25. For bt-CID experiments, HCD fragmentation was performed at a NCE of either 24, 27, 30, or 33%. For CID and HCD DDA experiments, fragments were analyzed in the Orbitrap at a resolution of 15,000 (200 m/z) with an AGC target of 200% and a max injection time of 22 ms.

For HEK 293T (Lumos) DIA experiments, MS1 scans of peptide precursors were performed at 120,000 resolution (200 m/z) over a scan range of 350-1050 m/z, with an AGC target of 250% and a max injection time of 100 ms. DIA scans were collected using 20 m/z staggered windows, with loop count set to 20, Orbitrap resolution = 15K, AGC target of 100%, and maximum injection time of 22 ms.

For HSC6 and SCC25 DDA experiments (Eclipse), eluting peptide cations were analyzed by electrospray ionization on an Orbitrap Eclipse (Thermo Fisher Scientific). MS1 scans of peptide precursors were performed at 120,000 resolution (200 m/z) over a scan range of 350-1050 m/z, with an AGC target of 250% and a max injection time of 100 ms. MIPS was set to peptide mode, and charge states 2-6 were selected for fragmentation. Dynamic exclusion was set to 20 s with a 10 ppm tolerance around the precursor and isotopes. The instrument was run in Top Speed Mode with a 3 second setting. Tandem MS was performed via quadrupole isolation with a width of 1.4 Th. For re-CID experiments, CID fragmentation was performed at a NCE of 27% or 30%, CID activation time of 10 ms, and activation Q of 0.25. For bt-CID experiments, HCD fragmentation was performed at a NCE of 27% or 30%. For CID and HCD DDA experiments, fragments were analyzed in the Orbitrap at a resolution of 15,000 (200 m/z) with an AGC target of 200% and a max injection time of 22 ms.

For HSC6 and SCC25 DIA experiments (Eclipse), MS1 scans of peptide precursors were collected as selected ion monitoring (SIM) scans performed at 120,000 resolution (200 m/z) over a scan range of 350-1050 m/z, with an AGC target of 200% and a max injection time of 100 ms. For re-CID experiments, CID fragmentation was performed at a NCE of 30%, CID activation time of 10 ms, and activation Q of 0.25, max injection time of 22 ms, and an Orbitrap resolution of 15,000 (200 m/z). For bt-CID experiments, HCD fragmentation was performed at a NCE of 30% max injection time of 22 ms, and an Orbitrap resolution of 15,000 (200 m/z). DIA scans were collected using 24 m/z staggered windows, with loop control set to All.

#### Database searching

Raw mass spectrometry data from each DDA dataset was independently searched using the Andromeda search engine^45^ built into MaxQuant^5^ (version 1.6.10.43). Samples were searched against a database of *homo sapiens* protein sequences (downloaded from Uniprot February 3, 2020). Each dataset was searched using default settings with the appropriate protease (either AspN, GluC, or trypsin) with up to two missed cleavages for the HEK 293T datasets and up to three missed cleavages for the HSC6 and SCC25 datasets. Carbamidomethylation of cysteines was set as a fixed modification, and oxidation of methionines and N-terminal acetylation were set as variable modifications. Peptides and protein groups were filtered to a 1% false discovery rate (FDR). Protein groups were filtered for “Only identified by site”, “Reverse”, and “Contaminant”. Spectral libraries for DIA searches were built from the MaxQuant msms.txt file using Spectronaut’s Pulsar search engine.^36^ Spectral libraries from all digestion protease were combined into a single library for each fragmentation mode and NCE. DIA data from each protease and protease mixture was searched independently using the corresponding spectral library and DIA-NN (version 1.7.16)^37^ For DIA-NN searches, raw data files were deconvoluted and converted to mzML files using ProteoWizard,^46^ with filter settings peakPicking (vendor msLevel=1-), demultiplex (optimization=overlap_only massError=10.0ppm), and titleMaker, with SRM as spectra selected for Eclipse experiments. In DIA-NN, MS1 mass accuracy was set to 5 ppm (suggested setting), and the neural network classifier was set to double-pass mode. Precursor FDR was set to 1%. Protein inference was performed at the Gene level using proteotypic peptides. Likely interferences were removed.

#### Data analysis

Protein concentrations were determined from collected trypsin digested DDA data using the Proteomic Ruler^39^ plug-in as part of the Perseus data analysis suite.^47^ Following DIA-NN analysis, protein group information was median normalized and log2 transformed. CVs were calculated on the non-log transformed replicate data and defined as σ/µ * 100%. Based on peptide quantifications reported by DIA-NN, protein level fold-changes and probability of differential abundance between biological groups were calculated using the Diffacto algorithm with default settings.^40^ Figures were generated in R version 4.0.3 (https://www.r-project.org) or InstantClue.^48^

## Results and Discussion

### Evaluation of DIA acquisition parameters for individual proteases

DIA proteomics workflows have been highly optimized for experiments utilizing tryptic peptides. Therefore, we began by evaluating data acquisition parameters for non-tryptic peptides. Here, we focused on fragmentation mode and collision energy, as we hypothesized these are parameters most likely to influence peptide identifications and spectral quality for peptides of varying chemistries. To determine the optimal settings for multi-protease DIA, DDA and DIA data was collected for three proteolytic digests (trypsin, GluC, and AspN) of HEK 293T cells at normalized collision energies (NCE) of 24, 27, 30, and 33 over an 80-minute LC-MS/MS gradient on an Orbitrap Fusion Lumos Tribrid mass spectrometer, with fragmentation by either bt-CID or re-CID.

The performance of DIA analysis is dependent upon MS/MS scan speed, relying heavily on the rapid scan speed (e.g. time-of-flight detectors) and bt-CID. Consequently, analysis of DIA data has primarily utilized y-ions,^30^ which are the predominant ion type generated from tryptic peptides upon bt-CID.^31^ However, due to the generation of peptides that typically lack a basic amino acid at the c-terminus, alternative proteases often do not generate a robust y-ion series. Thus we compared the average number of detectable fragment ions and their relative proportion between b- and y-type ions. We observed that under optimal collision energies the total number of fragment ions detected is similar across both fragmentation mode and protease. As expected, evaluation of the proportion of b- and y-type ions generated from tryptic peptides fragmented with bt-CID indicated that y-ions were favored,^32,33^ with the proportion of y-ions dramatically increasing at higher NCEs (**Fig. 1A**). In contrast, following fragmentation with re-CID, tryptic peptides produce nearly equal proportions of b-type and y-type fragment ions at all NCE values. Unlike trypsin, we find that for AspN and GluC, the proportion of b- and y-type ions is not dependent upon fragmentation mode, with nearly equivalent proportions being detected. Instead, these alternative proteases were more dependent upon NCE, with a low NCE of 24 resulting in a marked decrease in the number of fragment ions detected. (**Fig. 1A**). Thus, while all proteases detect similar numbers of fragment ions that can be utilized for peptide detection and quantification, the utilization of b-type ions in DIA analysis is an important feature for experiments utilizing either re-CID or alternative proteases.

**Figure 1.**
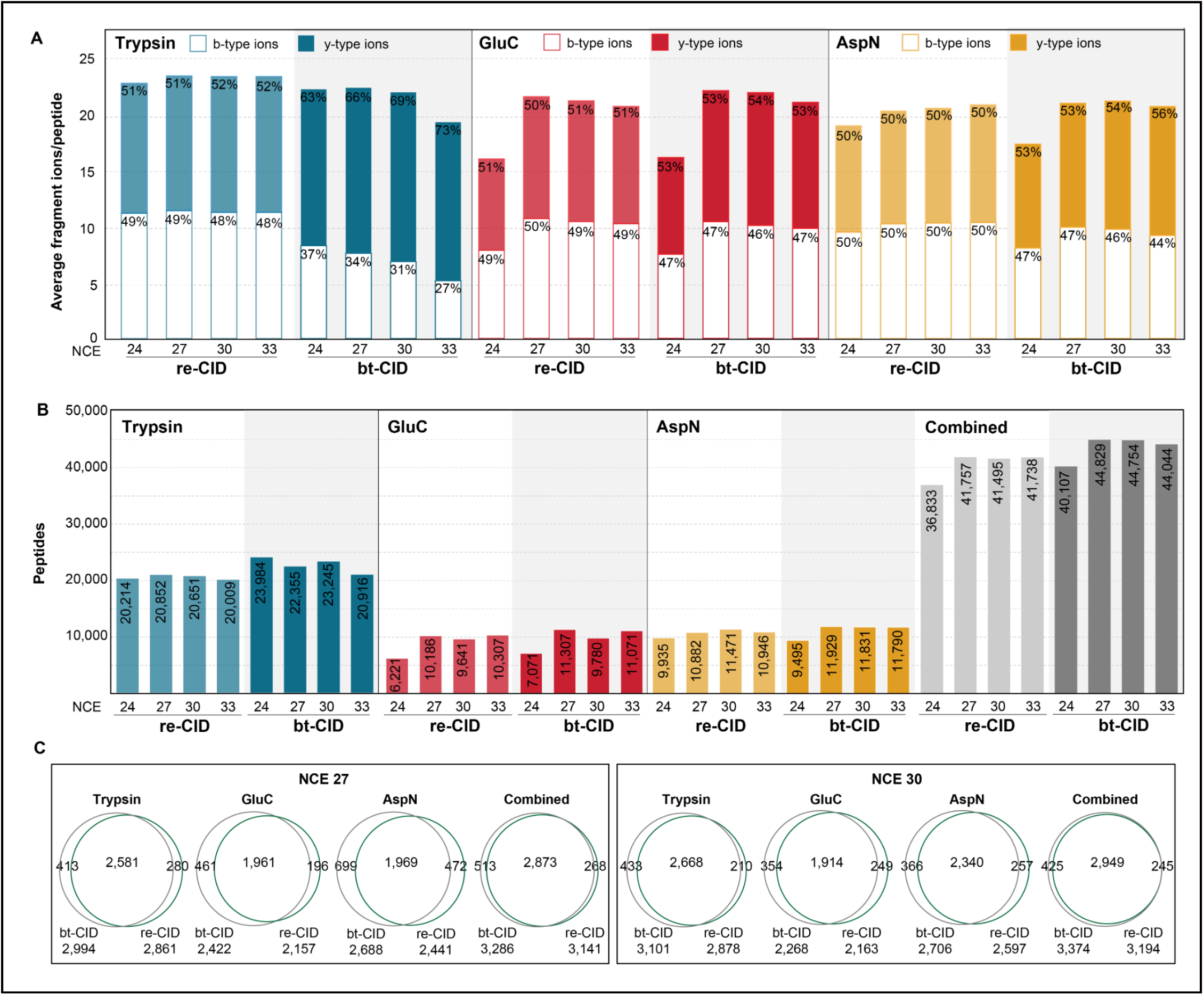
Evaluation of DIA acquisition parameters for individual proteases. (**A**) Average number of y-type (filled bars) and b-type (open bars) ions identified per protein in each spectral library. Percentages correspond to the proportion of y-or b-type ions in each spectral library. (**B**) Peptides identified in DDA experiments (colored bars) for each protease with the specified fragmentation and normalized collision energy (NCE). Combined peptides for all proteases at the specified fragmentation and NCE are plotted in grey. (**C**) Overlap between re-CID and bt-CID protein group identifications following DDA analysis for each protease at NCEs of 27 and 30.

Next, we evaluated the number of peptide detections and observed that bt-CID results in an average 8-10% greater peptide and protein identifications for nearly all NCE values evaluated, likely due to the moderately faster fragmentation of bt-CID (**Fig. 1B**). As expected, trypsin resulted in the detection of the greatest number of peptides,^20,34,35^ approximately twice that of GluC or AspN for both fragmentation modes. While the number of tryptic peptides identified was not impacted across the NCE values tested here, we observe a noticeable decrease in identifications for GluC and AspN at an NCE of 24, correlating with the reduced number of fragment ions detected at this collision energy.

We also compared the overlap in protein group identifications following DDA analysis for individual enzymatic digests following fragmentation at NCE of 27 or 30 with either re-CID or bt-CID (**Fig. 1C**). For all digests, we observed high overlap between both fragmentation methods, suggesting the validity of either method for subsequent DIA experiments. Based on these results, an NCE of 27 or 30 was selected as the optimal consensus NCE for both bt-CID and re-CID for subsequent DIA experiments.

### Increased peptide detections upon protease multiplexing DIA

Recent work has shown DIA-multiplexing of samples from different proteases can be used to significantly increase phosphopeptide detections when using bt-CID.^25^ Thus, we sought to determine if such benefits also extended to the analysis of unmodified samples. For this analysis, we used Spectronaut^36^ to combine the spectral libraries from duplicate 80 minute LC-MS/MS analyses of three individual proteases (trypsin, GluC, and AspN) into a single spectral library based on fragmentation and collision energy (**Supplemental Table 1**). DIA analysis was performed on either tryptic digests, or mixtures in which samples from individual proteases were pooled for DIA, and all DIA data was analyzed with DIA-NN^37^ (**Fig. 2**). Pooled samples were mixed in different proportions and each mixture was analyzed by DIA at two collision energies (NCE 27 and 30). The DIA results of this pooled sample were compared to DIA of trypsin alone. Here we find that DIA of pooled samples detected more peptides than trypsin alone in all conditions (**Fig. 3A**). When evaluating pooled samples mixed in differing proportions we find that the most peptides were detected when mixing protease in equal ratios (i.e. 1:1:1), because as tryptic peptides make up a smaller proportion of the mixture, both tryptic peptide detections and total peptide detections decrease. These decreases in tryptic peptide detections are compensated by only modest increases in peptides detected for GluC and AspN, despite peptides from these proteases being in 2-3 fold excess as tryptic peptides (**Fig. 3A**).

**Figure 2.**
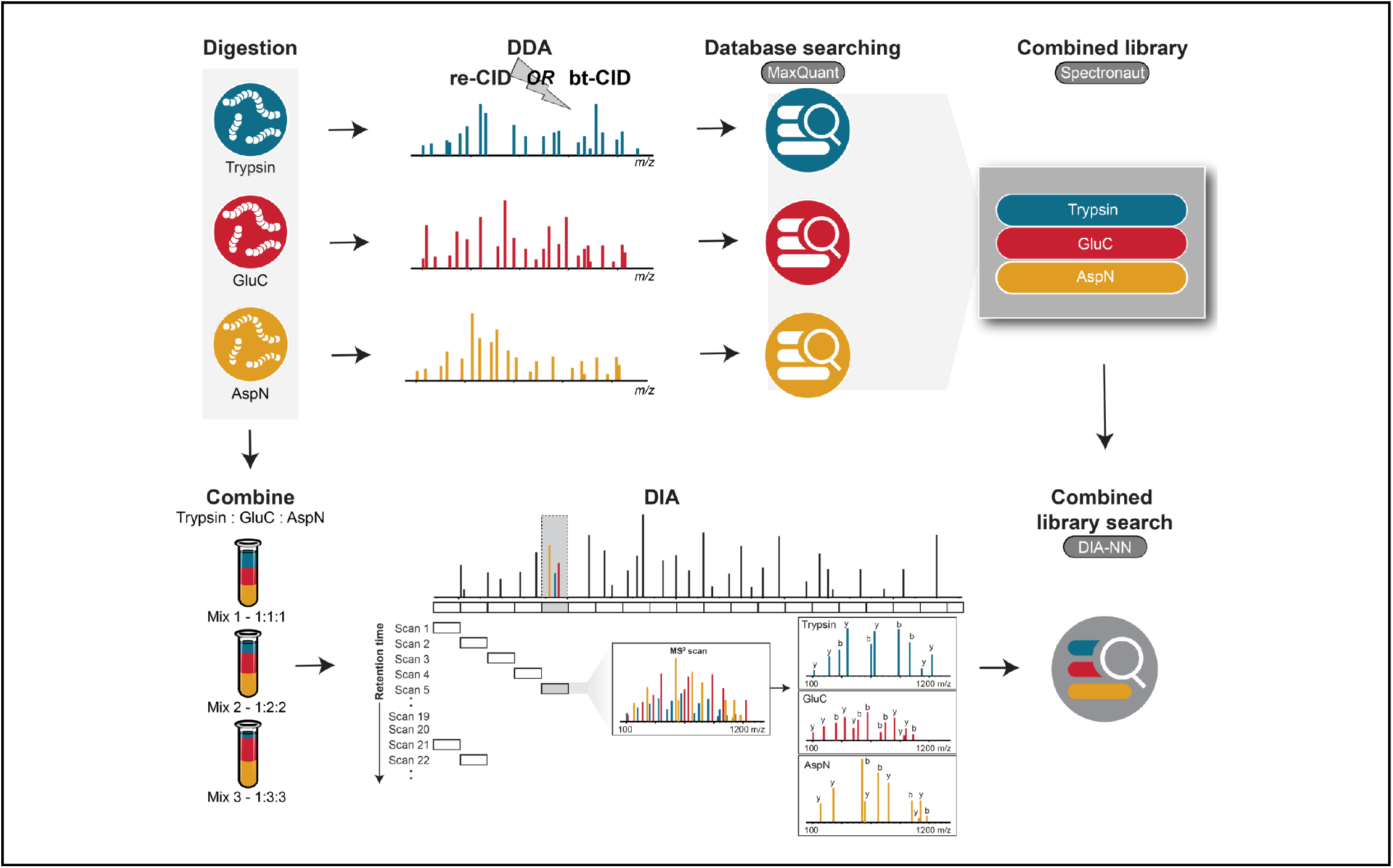
Overview of the DIA multiplexing strategy employed. DDA spectral libraries generated from samples individually digested with trypsin, GluC, or AspN were combined into a single library. Peptides derived from these three proteases were then mixed in varying proportions and the pooled sample was analyzed via DIA.

**Figure 3.**
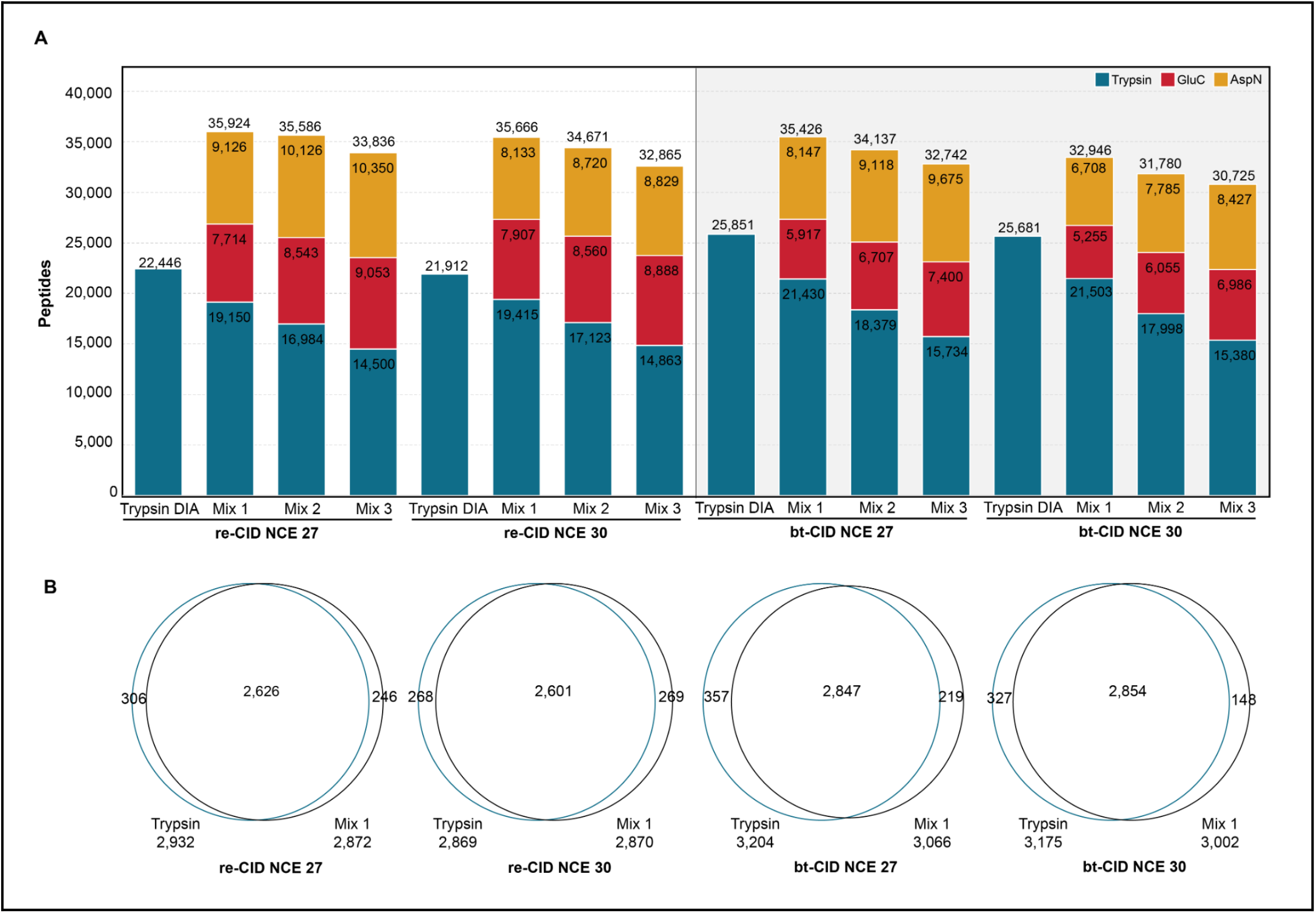
Peptide detection and recovery from multiplexed DIA samples. (**A**) Peptide identifications for multiplexed samples are compared with peptide identifications from DIA trypsin (blue bar) for each fragmentation method and NCE. (**B**) Venn diagram of protein overlap between trypsin DIA and Mix1 DIA for bt-and re-CID at NCE = 27 or 30.

The percentage of identifications recovered from the combined spectral library was highest for re-CID DIA at an NCE of 30 (73%, **Supplemental Fig. 1A**). Peptide recovery rates were slightly lower with bt-CID, identifying 61% and 63% of library peptides at NCEs of 30 and 27, respectively (**Supplemental Fig. 1A**). As the re-CID spectral libraries contain fewer peptides than the bt-CID libraries (**Supplemental Table 1**), total peptide identifications were similar between both fragmentation methods. Reproducibility at the peptide level between replicate samples was high for all fragmentation and NCEs, with at least 82% of peptides shared in duplicate runs for all analyses (**Supplemental Figure 1B**). Importantly, we observe that these significant increases in peptide identifications upon protease mixing in DIA are not detrimental to protein identifications, as nearly equivalent numbers of proteins are detected in the pooled samples as compared to trypsin alone (**Fig. 3B**). To test the validity of searching spectral libraries combined from different proteases, each dataset was searched with DIA-NN using a spectral library likely to contain only false positive matches. For example, trypsin-derived peptide samples were searched against a spectral library containing only AspN generated peptides. In all cases, DIA-NN could not identify enough matches to complete the searches, suggesting we are returning a low number of false positive identifications.

### Increased proteome sequence coverage upon protease multiplexing DIA

The increase in peptide detections upon protease multiplexing yields significantly greater protein sequence coverage compared to trypsin alone (**Fig. 4A)**. This increase is most notable for re-CID at NCE 27, where with trypsin we detect 19.2% of amino acids in the proteome, and upon multiplexing this increases to 27.9%, equating to a 27% relative increase in non-redundant amino acid residue detections compared to trypsin alone. For bt-CID, we observe a ∼4% increase in median proteome sequence coverage, equating to a 37% relative increase in non-redundant amino acid in multiplexed samples as compared to trypsin alone. However, bt-CID provided increased sequence coverage in more protein groups than re-CID. Specifically, following bt-CID analysis of the pooled sample (NCE 27), proteome sequence coverage increased for 1,441 protein groups by an average of 11.9%, while 1,086 protein groups had an average decrease in sequence coverage of 5.1% compared to trypsin alone. Overall, 337 proteins showed no change in sequence coverage. For re-CID (NCE 30), 1,798 protein groups saw an increase in proteome sequence coverage averaging 13.4%, while 533 protein groups had decreased coverage by an average of 4.0%. In this case, 297 protein groups had no change in sequence coverage. For each analyses, the distribution of the change in protein sequence coverage is plotted in **Supplemental Figure 2A**. The relationship between sequence coverage of individual protein groups and pooled digests is plotted in **Fig. 4B and C**. Notably, a number of protein groups were identified with 100% sequence coverage following multiplexed protease digestion, while none were identified with trypsin alone. The overlap in individual amino acid residues identified from either trypsin alone or the pooled samples are plotted in **Fig. 4D and F**. Using re-CID at NCE 30 (**Fig. 4D**), 412,031 unique amino acid residues were detected in the combined datasets, 29.9% (123,171) of which were not detected using trypsin alone. We found that bt-CID at NCE 27 identifies slightly more unique amino acid residues (480,719) across both datasets (**Fig. 4F**), with 37.7% (210,714) amino acids unique to the pooled sample. The contribution of these unique amino acid residues present in the mixed data is illustrated for SERF2 (P84101), a 50 amino acid protein containing many potential tryptic cleavage sites that render the majority of tryptic peptides too short to yield useful information. bt-CID analysis with trypsin identified a single tryptic peptide (**Fig. 2F**), while analysis of the multiple protease mixture identified 4, improving SERF2 sequence coverage from 13.6% to 86.4%, respectively.

**Figure 4.**
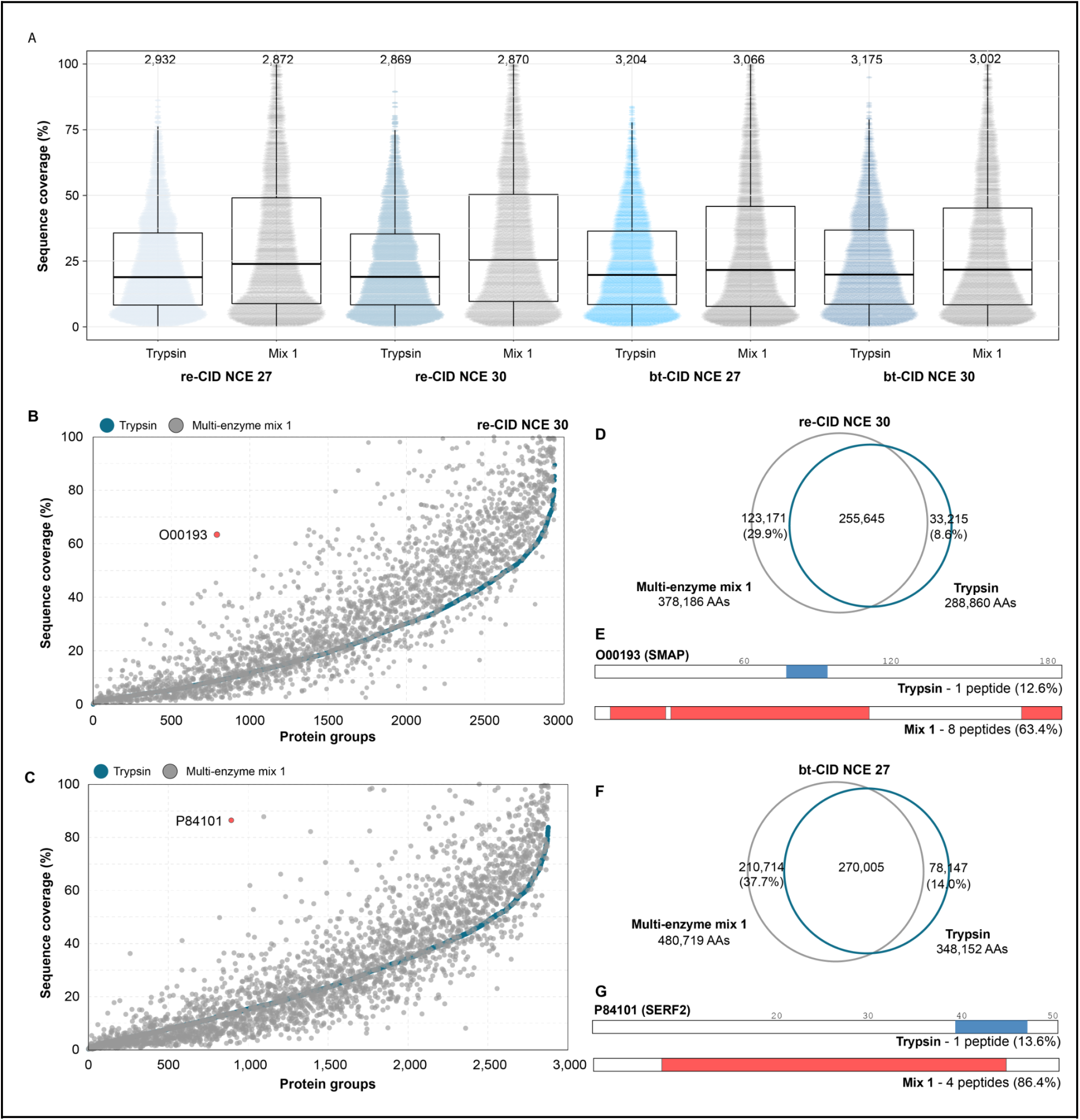
Sequence coverage. (**A**) Sequence coverage obtained from DIA analysis of trypsin (blue) and multiplexed mixture 1 (grey) for each fragmentation method and NCE. Each dot represents the sequence coverage for an individual protein group, with the total number of protein groups listed above. (**B**) Sequence coverage obtained for individual proteins following tryptic (blue) or multiplexed (grey) DIA analysis for re-CID at a NCE of 30 and (**C**) bt-CID at a NCE of 27 (bottom). (**D**) Overlap in observed amino acids following tryptic (blue) or multiplexed (grey) re-CID DIA analysis at a NCE of 30. (**E**) Schematic of sequence coverage of O00193 obtained using DIA re-CID and trypsin only (top, blue bar) or the multiplexed mixture (bottom, red bars). (**F**) Overlap in observed amino acids following tryptic (blue) or multiplexed (grey) bt-CID DIA analysis at a NCE of 27. (**G**) Schematic of sequence coverage of P84101 obtained using DIA bt-CID and trypsin only (top, blue bar) or the multiplexed mixture (bottom, red bars).

To determine the contribution of protein abundance on sequence coverage, we estimated the copies per cell for the protein groups in our dataset using the Proteomic Ruler method.^39^ For both fragmentation methods, proteins with the lowest expression levels showed little increase in sequence coverage (**Supplemental Fig. 2B**). Increases in sequence coverage correlated with protein abundance, with the largest gains in sequence coverage upon protease multiplexing seen for proteins present at over 100,000 copies per cell (**Supplemental Fig. 2B**). A similar trend is observed for the number of peptides per protein, which increased from 8 with trypsin alone to 14 with re-CID protease multiplexing (**Supplemental Fig. 2C**), or from 9 to 13 with bt-CID protease-multiplexing (**Supplemental Fig. 2D**). For both fragmentation methods, the largest increases in peptides per protein are in proteins with higher expression levels (**Supplemental Figs. 2E and F**). This suggests that the achievable depth of this method could be improved through incorporation of longer gradients or fractionation of samples prior to generation of spectral library. However, we do observe large sequence coverage increases for some proteins with low abundance. For example, SMAD (O00193) is estimated to be present in our dataset at ∼20,000 copies per cell. Only one tryptic peptide is identified by both re- and bt-CID, implying some potential utility for collecting mixed-mode or multi-injection re- and bt-CID measurements. Incorporation of other proteases increases the number of identified peptides to 8 with re-CID (**Fig. 4E, G**), boosting sequence coverage from 12.6% to 63.4%.

### Evaluation of quantitative performance

To assess the quantitative performance of our multiplexed protease DIA approach, we performed a comparative evaluation of two distinct head and neck squamous cell carcinoma cell lines. Equal aliquots of HSC6 and SCC25 cell lysate were individually digested with trypsin, AspN, or GluC. To build a combined spectral library, peptides from both cell lines were combined by protease and analyzed by re-or bt-CID DDA over a 120 minute LC-MS/MS gradient at a NCE of 30. For each cell line, the three distinct digests were then combined and analyzed in technical triplicate, using a 120 minute re- or bt-CID DIA method consisting of 24 *m*/*z* staggered windows. The results of these analyses were compared with trypsin digests of the same cell lines analyzed using the same DIA methods.

As observed previously, multiplexed DIA samples from multiple proteases (trypsin, GluC, and ApsN) resulted in significantly higher sequence coverage (**Supplemental Fig. 3A**). To determine the reproducibility and precision of our multiplexed protease approach, we calculated coefficients of variation (CV) for each fragmentation method. Each analysis was searched individually with DIA-NN against its matching combined database, and quantitative values were compared across replicates at the protein level. Median CVs for the multiplexed samples were similar across cell lines and fragmentation methods -with re-CID, we achieve CVs of 13.2% for HSC6 cells and 12.6% with SCC25 cells, and using bt-CID, CVs are 12.7% for HSC6 cells and 13.0% for SCC25 cells (**Fig. 5A**). These values are similar to the median CVs achieved with trypsin alone (∼ 11.1% with re-CID and ∼10.5% with bt-CID), but the multiplexed samples show a larger population spread in the third quartile. This may be linked to the combining of different peptide types. As tryptic peptides ionizes better than peptides derived from digestion with other proteases, we expect some variations when combining these peptides at the protein level. For both fragmentation methods, comparisons at the peptide level show similar CVs for the tryptic and non-tryptic-peptides identified in the mixture (**Supplemental Fig. 3B and C**).

**Figure 5.**
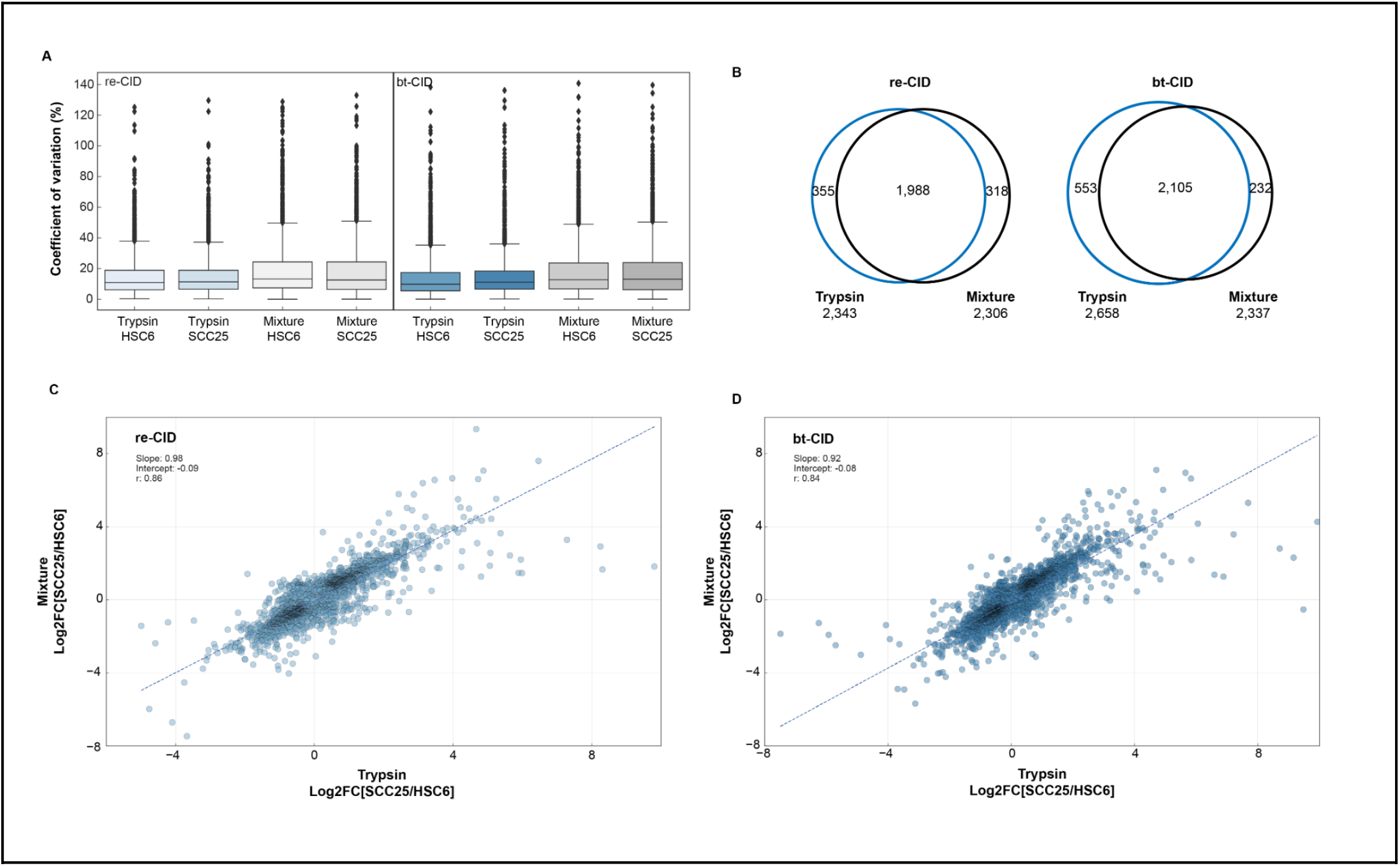
Evaluation of quantitative performance. (**A**) CVs for trypsin and mixture analyses of HSC6 and SCC25 cell lines using either re-CID or bt-CID. (**B**) Overlap of significantly changing proteins (corrected P-value <0.05) for trypsin and mixture analyses using either re-CID or bt-CID. (**C**) Correlation of log2 fold-change values (SCC25 vs. HSC6) for common proteins in the trypsin-only (x-axis) and mixture (y-axis) datasets analysed with re-CID. (**D**) Correlation of log2 fold-change values (SCC25 vs. HSC6) for common proteins in the trypsin-only (x-axis) and mixture (y-axis) datasets analysed with bt-CID.

Similar to our previous analyses, we observe an increase in the number of peptides identified following DIA analysis with the multiplexed mixture compared to trypsin alone (**Supplemental Fig. 3D**) -an increase to 51,207 from 27,316 peptides across the re-CID experiments and to 50,572 from 31,169 peptides across the bt-CID experiments. Sequence coverage of protein groups in the multiplexed mixtures also increased compared to samples digested exclusively with trypsin (Supplemental **Fig. 3A**), with re-CID showing the greatest boost in sequence coverage. Here, we also see a moderate increase in identified protein groups in the multiplexed experiments, increasing from 3,224 to 3,522 across cell lines and replicates with multiplexed re-CID and from 3,617 to 3,723 with multiplexed bt-CID. This increase may be attributable to the longer gradient time used in this set of experiments (120 minutes versus 80 minutes). We also observe a boost in the number of quantifiable peptides per protein in the multiplexed mixtures, increasing from a mean of 7.9 peptides per protein in the trypsin-only sample to 12.6 peptides per protein with multiplexed re-CID, and an increase from 8.6 to 11.2 quantified peptides per proteins in the multiplexed bt-CID samples (**Supplemental Fig. 3E and F**).

We next compared the relative quantitative accuracy of proteins identified by either trypsin alone or with our multiplexed approach. To control for possible discrepancies in protein-level quantification potentially arising from covariances in peptide abundances, we used Diffacto^40^ to perform relative quantitative analyses between the HSC6 and SCC25 cell lines. To assess the number of protein groups significantly changing between the two cell lines, we focused on the corrected p-values (PECA score) calculated for the trypsin-only and mixture datasets for both fragmentation methods. Analysis of the re-CID datasets shows a similar number of significantly changing proteins (corrected p-value < 0.05) in the trypsin (2,343) and mixture (2,306) samples. Of those proteins, 1,998 were common between both datasets, with 355 significantly changing proteins unique to the trypsin dataset and 318 significantly changing proteins unique to the mixture (**Fig. 5B**). The overlap of proteins was also high for the bt-CID dataset at 2,105 proteins (**Fig. 5B**), although more significantly changing proteins were quantified in the trypsin-only dataset (2,658) than the mixture (2,337).

We next compared the similarities of fold changes between the two cell lines in the trypsin-only and mixture datasets. A correlation plot of the log2 transformed fold-change values (SCC25 *vs* HSC6) of the 3,042 proteins common to both datasets shows a strong correlation between samples (Spearman’s R = 0.86) for the re-CID data. Similar correlation values (Spearman’s R = 0.84) are observed for the 3,178 proteins common to the trypsin and mixture datasets following bt-CID analysis. Combined, these results indicate that protease multiplexing can provide similar quantitative performance as trypsin alone, with the benefit of increased peptide identifications and sequence coverage.

## Conclusions

Here we present a readily accessible multiplexed DIA workflow for the analysis of several distinct proteolytic digests within a single sample, and provide guidance for developing re-CID based DIA experiments for tryptic and non-tryptic peptides. We demonstrate the utility of this method for both re-and bt-CID to increase identified peptides and sequence coverage, with minimal increases MS acquisition times, and without compromising achievable proteomic depth. The increase in amino acid residue coverage gained with this methodology may be helpful in distinguishing between expressed proteoforms^41,42^ that can require near complete sequence coverage for assignment. Potential limitations of this methodology are that the largest gains in sequence coverage are often from high abundance proteins, and that the increased complexity of multiplexed samples can impact quantitative precision. However, these limitations can likely be overcome through the incorporation of longer LC-MS/MS gradients, or pre-fractionation of samples prior to generation of spectral libraries. While we typically detected fewer peptides with re-CID than with bt-CID in this experiment, this is likely due to smaller library sizes resulting from slower DDA MS/MS rates with re-CID. In this case, newer library generation methods to predict re-CID fragmentation^43^ or the conversion of bt-CID libraries to re-CID^44^ may improve detection results when analyzing re-CID datasets with DIA. Considering the relative immaturity of DIA analysis, particularly for non-tryptic peptides, we anticipate that further work will illuminate what increase in proteome coverage can be routinely observed, as well as which sample types and biological questions are most benefited by this strategy, such as the previously demonstrated application of phosphorylation site analysis.^25^

## Acknowledgements

We thank Nevan J. Krogan for use of the Thermo Fisher Scientific Proteomics Facility for Disease Target Discovery at the Gladstone Institutes, and the lab of Jack Taunton for the use of their Thermo Scientific Orbitrap Eclipse Tribrid mass spectrometer for data presented in Figure 5.

## Funding

NIH R01GM133981 to BCS and DLS. DBW is additionally supported by NIH K99HD090201.

## Contributions

ALR, BCS, and DLS conceived and designed the project. KC, ES, and ALR performed the experiments. ALR and DBW performed the data analysis, ALR, DBW, BCS, and DLS drafted and revised the manuscript which has been seen and approved by all authors. BCS and DLS supervised the work.

## Competing interests

BCS is a founder and shareholder in Proteome Software, which operates in the field of proteomics. DLS has a consulting agreement with Maze Therapeutics.

## Data availability

All raw MS data files, search results and individual spectral libraries are available from the Pride partner ProteomeXchange repository under the PXD027242 identifier.

