## Supplementary Information for "Data-independent acquisition protease-multiplexing enables increased proteome sequence coverage across multiple fragmentation modes"

**Supplemental Table 1 –** Peptide identifications for individual spectral libraries and the merged spectral library. Individual libraries were constructed in Spectronaut from MaxQuant msms.txt files. Combined spectral libraries were constructed from individual libraries using the Merge feature in Spectronaut.

| | | Individual spectral libraries | | | Combined spectral library | |
|---|---|---|---|---|---|---|
| **Fragmentation** | **NCE** | **Tryptic peptides** | **GluC peptides** | **AspN peptides** | **Precursors** | **Combined peptides** |
| re-CID | 27 | 23,964 | 11,587 | 12,779 | 59,140 | 48,258 |
| re-CID | 30 | 23,310 | 11,628 | 12,157 | 57,061 | 47,022 |
| bt-CID | 27 | 28,195 | 13,578 | 12,063 | 65,005 | 53,769 |
| bt-CID | 30 | 28,262 | 13,212 | 14,503 | 69,138 | 55,902 |

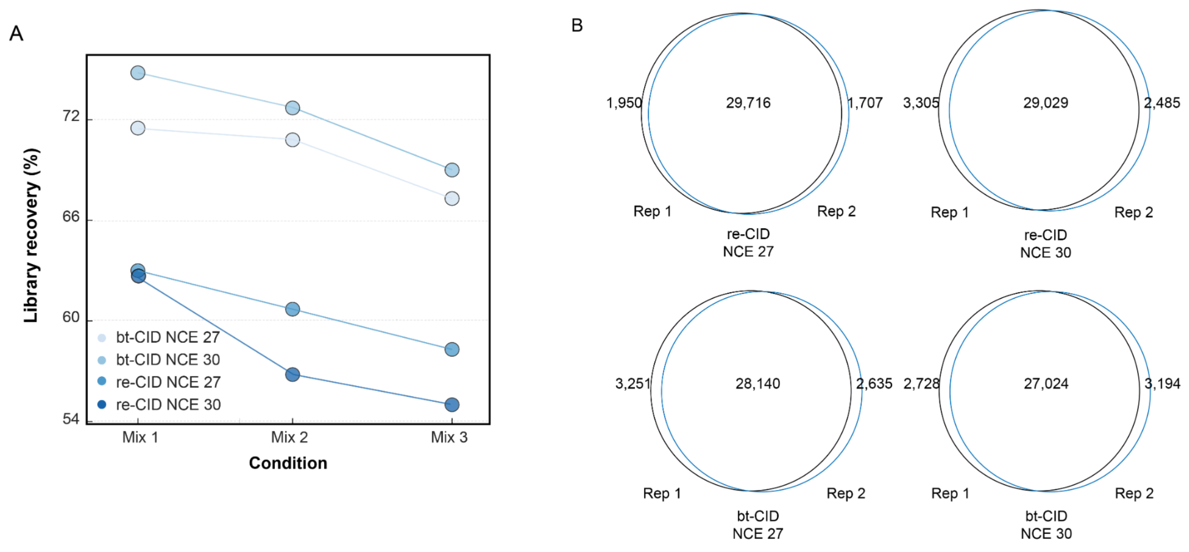


**Supplemental Figure 1**. (**A**) Percent of peptides in the spectral library identified in multiplexed DIA analyses. (**B**) Reproducibility between technical replicates at the peptide level for multiplexed analysis.

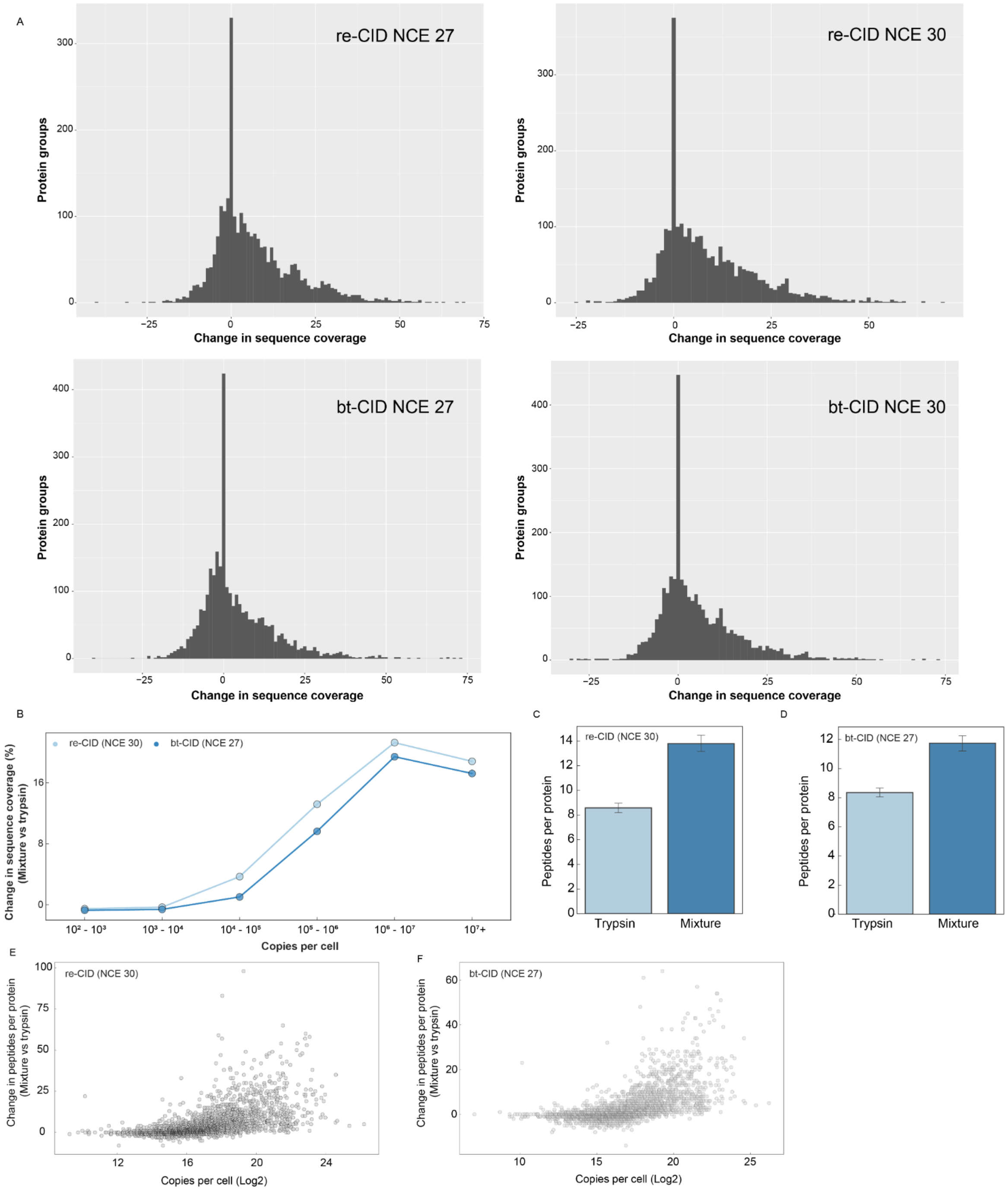


**Supplemental Figure 2**. (A) Histograms of distribution of change in sequence coverage between mixture and trypsin samples for each fragmentation energy and collision energy. (B) Change in sequence coverage as a function of protein abundance. Number of peptides identified per protein for trypsin (light blue) and mixture (blue) for re-CID (C) and bt-CID (D).

Number of peptides per protein as a function of protein abundance for re-CID (E) and bt-CID (F).

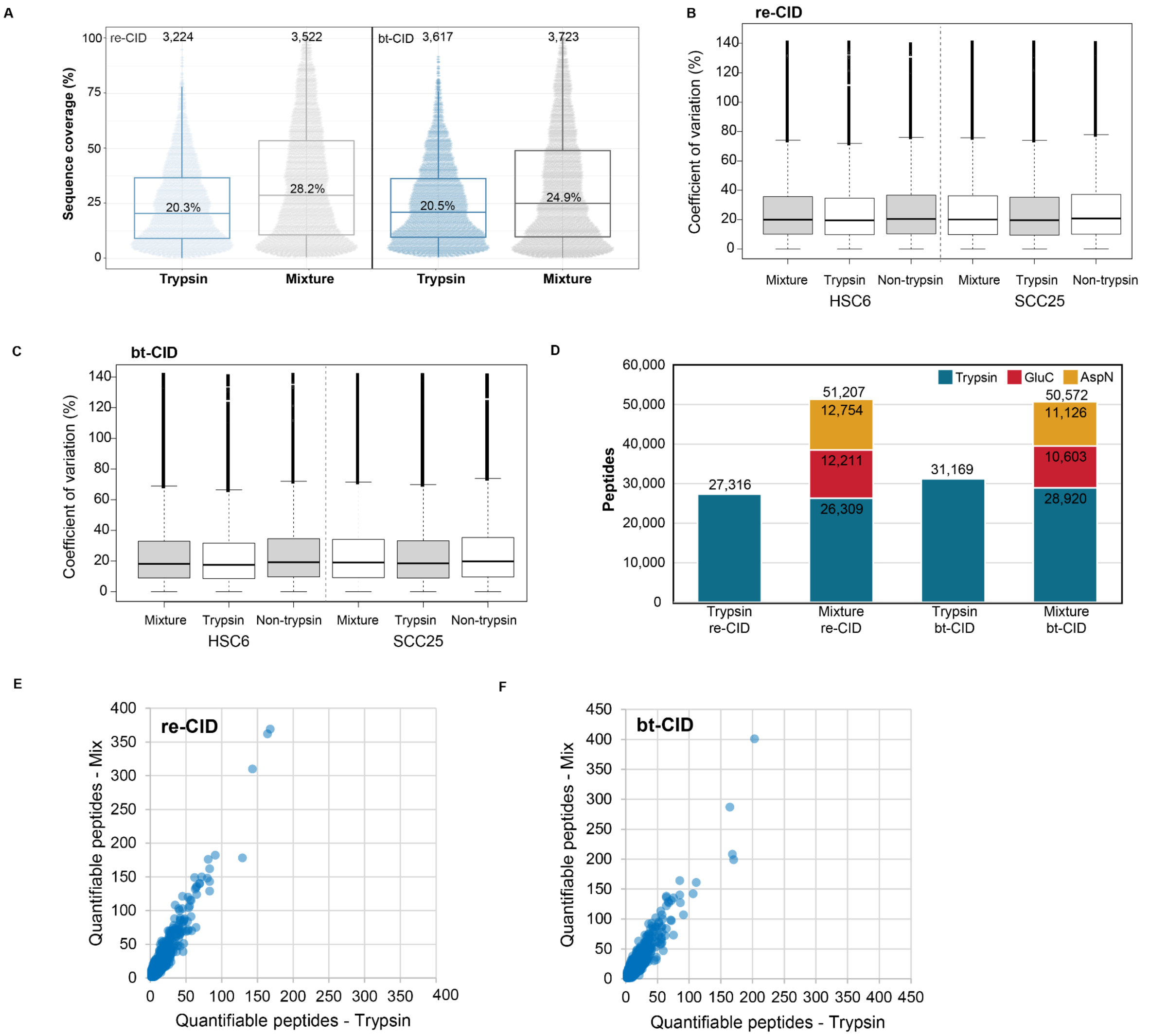


**Supplemental Figure 3.** (A) Boxplots of distribution of sequence coverage for proteins identified in trypsin-only (blue) and mixture samples (grey) in quantitative experiments analyzed with re-CID or bt-CID. (B) Peptide level CVs of tryptic, non-tryptic, and all peptides identified from the mixture samples (HSC6 and SCC25) following re-CID analysis. (C) Peptide level CVs of tryptic, non-tryptic, and all peptides identified from the mixture samples (HSC6 and SCC25) following bt-CID analysis. (D) Peptides per protein used for quantitation in trypsin-only (x-axis) and mixture (y-axis) samples analyzed with re-CID. Each dot in the plot represents a protein common to the two sample sets. (E) Peptides per protein used for quantitation in trypsin-only (x-axis) and mixture (y-axis) samples analyzed with bt-CID. Each dot in the plot represents a protein common to the two sample sets.
